# Application of Bayesian-Derived Gene Interaction Network and Subnetwork for Classification of Idiopathic Parkinson’s Disease with Biomarkers

**DOI:** 10.64898/2026.10.05.756343

**Authors:** Ruicheng Yang, Yaxia Yuan

## Abstract

Given the expenses associated with Parkinson’s Disease (PD), it is imperative to develop a method for accurate diagnosis before emergence of motor symptoms. While there are tests that can measure early onset of PD, these tests still rely on subjective input from patients and can thus be biased. Factors influencing their accuracy include environmental agents such as education and culture. Thus, blood-based transcriptomic signatures are a promising direction for biomarkers in PD that are minimally invasive and more objective than clinical evaluation. In this paper, we analyzed gene expression from the PPMI cohort. Using a Bayesian framework, a gene interaction network was constructed to enhance differential expressed genes (DEG) analysis through signal diffusion. In tandem with clinical traits, 13 genes were selected for optimal discrimination using a random forest model. Within the holdout test set, the model achieved an AUC of 0.972, outperforming contemporary models. The model’s performance demonstrates the potential of using blood-based transcriptomics to enhance clinical diagnosis of PD. Other genes recovered by network diffusion but missed by traditional DEG analysis are already validated as potential biomarkers in contemporary PD research. By combining transcriptomic signatures with clinical evaluation, accurate diagnosis of PD can happen before the major destruction of dopaminergic neurons, which is critical for effective intervention and treatment.

## Introduction

Parkinson’s disease (PD) is the second most prevalent neurodegenerative disorder. By 2037, an estimated 1.6 million people in the U.S. are projected to be affected by PD [1]. Thus, it is crucial to find a safe and effective tool for early detection of the condition. The current limitation of effective diagnosis is that identification often comes after the emergence of motor symptoms.

Recent studies suggest that by the time of PD onset, 60 to 80 percent of an individual’s dopaminergic content are already lost [2]. Furthermore, tests such as seed amplification assays (SAA) for alpha-synuclein achieve a high level of specificity and sensitivity, but often require invasive procedures due to needing samples from the cerebrospinal fluid (CSF) [3] [4] [5].

Currently, blood transcriptomics is becoming a promising direction for biomarker discovery in PD [6] [7] [8]. Furthermore, the Parkinson’s Progression Markers Initiative (PPMI) conducted a large-scale longitudinal study that measures gene expression through whole-blood RNA-sequencing [9]. This has become a widespread source for biomarker research. Recent studies have combined the expression of transcriptomic signatures with traits such as sex, Montreal Cognitive Assessment (MOCA) score, and University of Pennsylvania Smell Identification Test (UPSIT) scores [10]. While the UPSIT is linked to cognitive function, its assessment is still based on patient feedback and can be subject to inaccuracy [11]. Meanwhile, the MOCA test is more objective than a questionnaire like the Unified Parkinson’s Disease Rating Scale (UPDRS) Part 1 and also directly related to cognitive function. However, since factors like a patient’s native language or education level can influence results, MOCA alone is not a sufficient tool to diagnose PD either [12] [13]. In this regard, an advantage that blood transcriptomics offer compared to clinical evaluation is a purely objective measurement that can eliminate biases in diagnosis. Thus, models combining transcriptomics and clinical traits often achieve high specificity and sensitivity. Yet, many past panels of transcriptomic signatures achieve limited utility when applied to the PPMI cohort separate from clinical traits [14].

Furthermore, the discovery of biomarkers in patients with idiopathic PD is one of the most challenging aspects of proper diagnosis. This is due to the lack of a single genetic cause in individuals with idiopathic PD, leading to heterogeneity of symptoms in said patients [15] [16]. Finally, when doing gene expression analysis, many studies that focus on PD do not take into account gene interactions to enhance discovery of relevant genes. Methods like DiSNEP use correlation and validated interactions to achieve higher performance than traditional analysis [17].

In this study, we conducted a longitudinal analysis of all samples that were either idiopathic PD or healthy control. Using gene network diffusion to enhance differential expression gene (DEG) analysis, signals were chosen that represented their biological relevance regardless of time points. These transcriptomic signatures were analyzed for pathway enrichment and immune cell infiltration to ensure they represented biological relevance rather than noise. Relevant signatures were chosen to enhance the diagnostic accuracy of PD using only clinical evaluation. Finally, a classification model using 13 transcriptomic signatures and relevant patient traits was trained on baseline samples that outperformed models trained solely on clinical evaluation.

## Material and Methods

### Data Collection and Processing

All data for this study were collected from the PPMI RNA-Sequencing cohort via LONI IDA. Samples that were included for this analysis must have had the disease status of either healthy control or idiopathic PD. The PPMI dataset also provided information such as clinical visit and sex of each sample. Using the design developed by Xu et al. [18], sex, percent of mRNA bases, and age at consent squared were included as biological factors in differential expressed gene (DEG) analysis. Principal component analysis (PCA) was also conducted using the values already provided by the cohort. From exploratory PCA plots, any additional technical confounders that yielded in stratification or clusters were included as covariates DEG analysis.

### Differential Expression Analysis Baseline

All DEG analysis was performed in the limma-voom framework of 3.68.4 in R [19]. Genes with low expression counts across samples were automatically excluded in the pipeline. After concluding no technical factors that contributed to the variance of the top two principal components, analysis followed the final design = 0 + sex + percent mRNA bases + age squared + disease status. Corrected for multiple hypothesis testing, adjusted p-values were obtained using the Benjamini-Hochberg control method.

### DiSNEP Network-Diffusion Analysis

Only genes that had a log-2-fold change magnitude of greater than 0.1 were included in the network construction process. Following that, the correlation network was constructed based on the WGCNA package in R [20]. A soft threshold power of 27 was implemented for topological overlap matrix (TOM). All edges in the TOM with a weight greater than 0.15 were transformed into values of 1, while all remaining edges became 0 in a binary correlation matrix. Only the top 10% of edges in the correlation matrix were kept the same. For enhanced network diffusion, a STRING network was also built with the threshold of confidence for interactions between two genes being set at 1. Finally, after normalization of the correlation matrix, the DiSNEP framework developed by Ruan et al. [17] was implemented. A seed of 67 was set in R.

Following the workflow, each gene’s adjusted p-value from traditional DEG analysis were transformed into association signals based on the expression *g* = ϕ(1 − *p*). The enhanced gene network was used for a similar random walk process shown below,

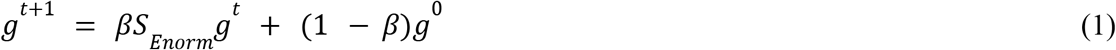

where *β* was assigned a value of 0.75 for optimization, S was the correlation enhanced STRING network, and g was the vector of association signals for genes that passed the log-fold change threshold. Once the enhanced association signals were obtained, they were transformed back to their respective p-values. Finally, if a gene passed the log fold change and transformer p-value thresholds of 0.1 and 0.05, respectively, they were deemed to be significant DEGs.

### Bayesian Network Analysis

Building on top of the framework proposed by Huang et al. [21], an adjacency matrix and correlation matrix were implemented for a hyperparameter π based on the following equation,

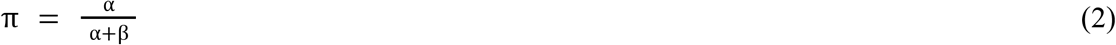

where α was assigned a value of 10 if the edge entry between two genes in the STRING adjacency matrix was non-zero, otherwise assigned a value of 1. Like DiSNEP, only genes that had a log-fold change magnitude of greater than 0.1 were included in the network. A similar approach was taken for β, only a value of 5 was assigned if the edge entry between two genes in the binary correlation matrix was non-zero. This was done to ensure that gene-interactions validated in literature had more importance than interactions from a correlation network.

The probability of an interaction existing between two genes p was assumed to follow a Bernoulli distribution based on the hyperparameter π. The third hierarchy focuses on the strength of an interaction between two genes given that the interaction exists and follows the distributions provided by Huang et al [21]. Lastly, given the gene interaction strength, the normalized expression of a gene in a sample is assumed to follow a normal distribution centered on the sum of the product of every other gene’s expression times the strength of the given interaction with the gene of interest. Metropolis-Hastings algorithm was implemented for network construction using 500 iterations, with a set seed of 67 in R. The equation for this distribution is shown below, with variance assumed to be 1. All interactions with a probability of existence greater than 0.5 were assumed to be existing interactions and were transformed into entries of 1, with all other entries being 0. This matrix was then normalized for the same signal diffusion process as DiSNEP for recovery of genes.

### Bayesian Subnetwork Analysis

Following the Bayesian network construction, a novel method for subnetwork analysis was developed to identify gene modules. Let γ denote the edge existence interaction matrix and *Θ* denote the biochemical interaction-strength matrix obtained from the previous Bayesian model. A disease-specific weighted interaction network was constructed as *A* = γ ° *Θ*, where ° denotes the Hadamard (element-wise) product. Then, only genes with a p-value of less than 0.1 or belonging to the top 15% highest-degree hub genes were kept as nodes in the disease-specific network. For computational efficiency, every patient was assumed to have the same topological gene interaction network, where only the node features–represented by gene expression–varied between samples.

A seed of 67 was set in R to run both the graph convolutional network (GCN) and graph attention network (GAT) algorithms. For the GCN layer, suppose every patient can be written as a node feature matrix *H*^0^ = [*X*_1_, *X*_2_, … *X*_*i*_]^*T*^, where *X*_*i*_ is the normalized expression of a given gene i for the patient. However, since each gene currently only has 1 feature. Thus, at the beginning of training, a matrix *W* of dimensions 1×64 is initialized randomly once before training. For each patient, a new matrix is created using the linear projection method *Z* = *H*^0^ *W*_*GCN*_ so that every gene has 64 features. The adjacency matrix for the GCN layer is transformed using the equation, Â = *D*^−1/2^(*A* + *I*’)*D*^−1/2^, where A is the original weighted matrix, D is the degree matrix in which the diagonal entries are the row sums of *A* + *I*’, I’ is like the identity matrix–only the diagonal entries are the maximum weight of 0.9–and Â is the transformed interaction matrix. Finally, the hidden features matrix for each patient can be expressed in the equation *H*^(1)^ = *ReLU*(Â*Z*). *H*^(1)^ becomes a feature matrix that not only captures individual gene expression, but also gene relationships from the adjacency matrix.

The second half of the subnetwork analysis involves the implementation of a GAT layer that outputs the attention importance of each pair of genes. First, *H*^(1)^ is used in another pass of linear projection to form the new matrix *Z*^*^ = *H*^(1)^*W*_*GAT*_, where the *W*_*GAT*_ is again randomly initialized. Two learnable vectors *a*_*s*_ and *a*_*d*_ are also randomly initialized, used to evaluate every edge interaction in the network. Then, for any given interaction between gene i and gene j, ε_*ij*_ is equal to *LeakyReLU* 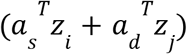 . Given that, the attention weight *α* for gene i and gene j is expressed in the formula 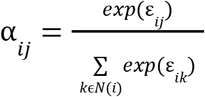, where k represents all the genes connected to i. Each gene is then transformed to a final hidden feature vector 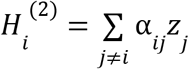 .

With both the GAT and GCN layers, a final GNN layer was constructed to classify a sample as either healthy control or idiopathic PD based on an output matrix ŷ = [*y*_1_, *y*_2_], where *y*_1_ is the probability of a patient being healthy and *y*_2_ is the probability of a patient being an idiopathic PD case. To do this, *H*^(2)^ is transformed into a single vector *R* per patient via. the formula 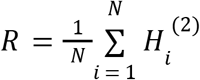 . In the final GNN layer, 2 weight matrices for two fully connected layers *W*_*FC*1_ and *W*_*FC*2_ are randomly initialized to execute the following 2 equations,

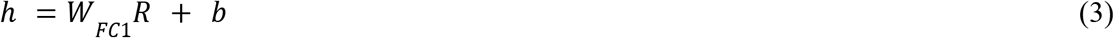

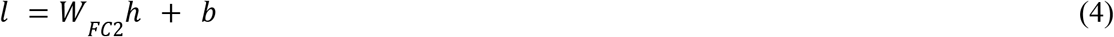

where *l* is expressed in the format *l* = [*l*_*healthy*_, *l*_*iPD*_]. It is important to note that a ReLU activation followed by dropout was used before the second fully connected layer. Using the logits, the predicted probability of a patient belonging to either class–represented by ŷ_*i*_ –is the final output. The classification loss L is based on the expression 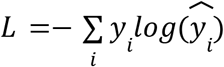, where y is the actual outcome in the format of a binary vector. For every iteration after the first iteration and up to 40, a random variable G will be updated using the gradient descent rule 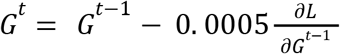 . This gradient descent rule was applied to all matrices and variables that were randomly initialized in both the GCN and GAT layers. The final attention matrices after 40 iterations were extracted with the batch loss converging at approximately 0.65 after 10 iterations.

If the average of the attention weights between gene i and j has a smaller magnitude for iPD patients compared to healthy controls, then the gene pair is determined to have a weaker interaction, classified as a “Loss of Function.” Conversely, a larger magnitude in average attention weights for iPD patients is assigned as a “Gain of Function” for the gene pair. Finally, Louvain clustering was used to cluster genes together into subnetwork modules. Any subnetwork with a size less than 5 was discarded.

To rank the genes in each module, each gene is given an eigenvector centrality *E*_*i*_ that gives a high score to genes that are connected to other highly connected genes. Each gene was also given a score of 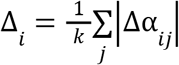, where k is the number of nodes connected to the gene of interest and Δα_*ij*_ is the change in attention for an interaction between case and control samples. Using the module expression matrix, PCA was also performed and each gene was assigned a score equivalent to their absolute loading on the first principal component *L*_*i*_ . Each metric was normalized to 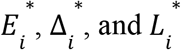 . Finally, each gene’s score was calculated using the equation ^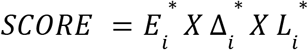^ . The top 10 scoring genes from each module were selected as significant DEGs.

### Machine Learning Sample Assignment

All the samples were divided with a training:testing split of 80:20. Stratified splitting was conducted to ensure no patient had multiple measurements in both the training and validation set. The library size of the validation set was determined through calcNormFactors in R and expression was normalized using log-2 counts-per-million (CPM), following the same assumptions as the training dataset normalization. The broader training set consisted of 1,695 samples and was used for DEG analysis, gene-network construction, and feature selection. This was done to ensure that the transcriptomic signatures chosen reflected reliable signals across the progression of PD using all time points and to filter out noise in the baseline samples. The broader testing set consisted of 448 samples.

### Feature Selection and Model Construction

LASSO regression was performed in R to simplify the model and prevent overfitting for expression matrices of significant genes from DiSNEP Network, Bayesian Network and Subnetwork analysis [22]. During LASSO feature selection, internal 10-fold cross validation was applied according to a stratified split so that a patient had all its samples in only one fold. While the subnetwork analysis only yielded at most 5 transcriptomic features, the top 47 transcriptomic features were chosen from DiSNEP and Bayesian network DEGs after LASSO regression, respectively. This was determined due to the Bayesian network yielding 47 features after LASSO regression was applied. Genes from all 3 methods were pooled together. Network topological analysis was performed to identify all genes from the pool that had a connectivity of 5 edges or greater. From there, samples in both the training and testing sets were filtered out if they were not measured at baseline for model construction and evaluation. To prevent leakage, the baseline training and testing sets were normalized again using the same procedure only with a different background for the TMM method. Any sample with missing values for either UPSIT or MOCA were excluded. In the end, the baseline training and testing sets consisted of 400 and 102 samples, respectively. Using the hub genes in tandem with MOCA and UPSIT scores, a random forest model with 500 decision trees was trained on samples that were in the baseline training set.

In the event that the constructed model outperformed another random forest model trained only on MOCA and UPSIT scores, A SHAP matrix was also constructed to see which features pushed the recovered samples toward the correct prediction of idiopathic PD. Transcriptomic features were selected for consideration based on three criteria. The first was that the expression of the genes that pushed towards the prediction of PD must not contradict the general direction of the expression change for the genes in recovered PD samples compared to healthy control samples. The second criteria was that the average SHAP value had to be greater than a threshold of 0.001. This was determined by the MOCA score feature having an SHAP value of 0.0014. The final criterion was that the change in expression between recovered PD samples and healthy control samples must be consistent between the training and testing split for reliability. From there, the signatures that achieved the best AUC were considered for further analysis.

### Biological Characterization of DEGs

Gene Ontology (GO) Term enrichment was conducted on all significant DEGs uncovered by DiSNEP and Bayesian network/subnetwork analysis. Kyoto Encyclopedia of Genes and Genomes (KEGG) Term enrichment was also performed on the significant genes. The R packages used included clusterProfiler and enrichplot for visualization [23] [24]. Multiple hypothesis testing was performed using the Benjamini Hochberg procedure.

### Gene Set Variation Analysis (GSVA) enrichment

In a similar workflow to the one used by Wang et al. [25], the GSVA package in R [26] was adopted to compare pathways between high-and-low expressions of the final transcriptomic signatures. High and low expression was determined based on the cutoff expression settled at the median for each hub gene. For interpretability, the group with higher expression of a hub gene and a t>0 value for a pathway signifies that the pathway is more activated for said group. The group with lower expression of a hub gene and a t<0 value for a pathway signifies the same thing. Specifically, associations with an |*t*| greater than 1 were deemed significant.

### Immune-cell Infiltration

The “CIBERSORT” algorithm was used in R to identify differentially expressed immune cell types between idiopathic PD patients and healthy controls [27] [28]. Deconvolution utilized samples from all time points. In contrast to the classification model construction, immune cell type deconvolution utilized transcript-per-million (TPM) datasets that were also available from the PPMI cohort. However, only samples at the baseline measurement were used for further analysis after deconvolution. Finally, correlation analysis was performed to identify possible associations between the transcriptomic signatures and the cell types. The workflow of the model development and further analysis is shown in Figure 1.

**Figure 1.**
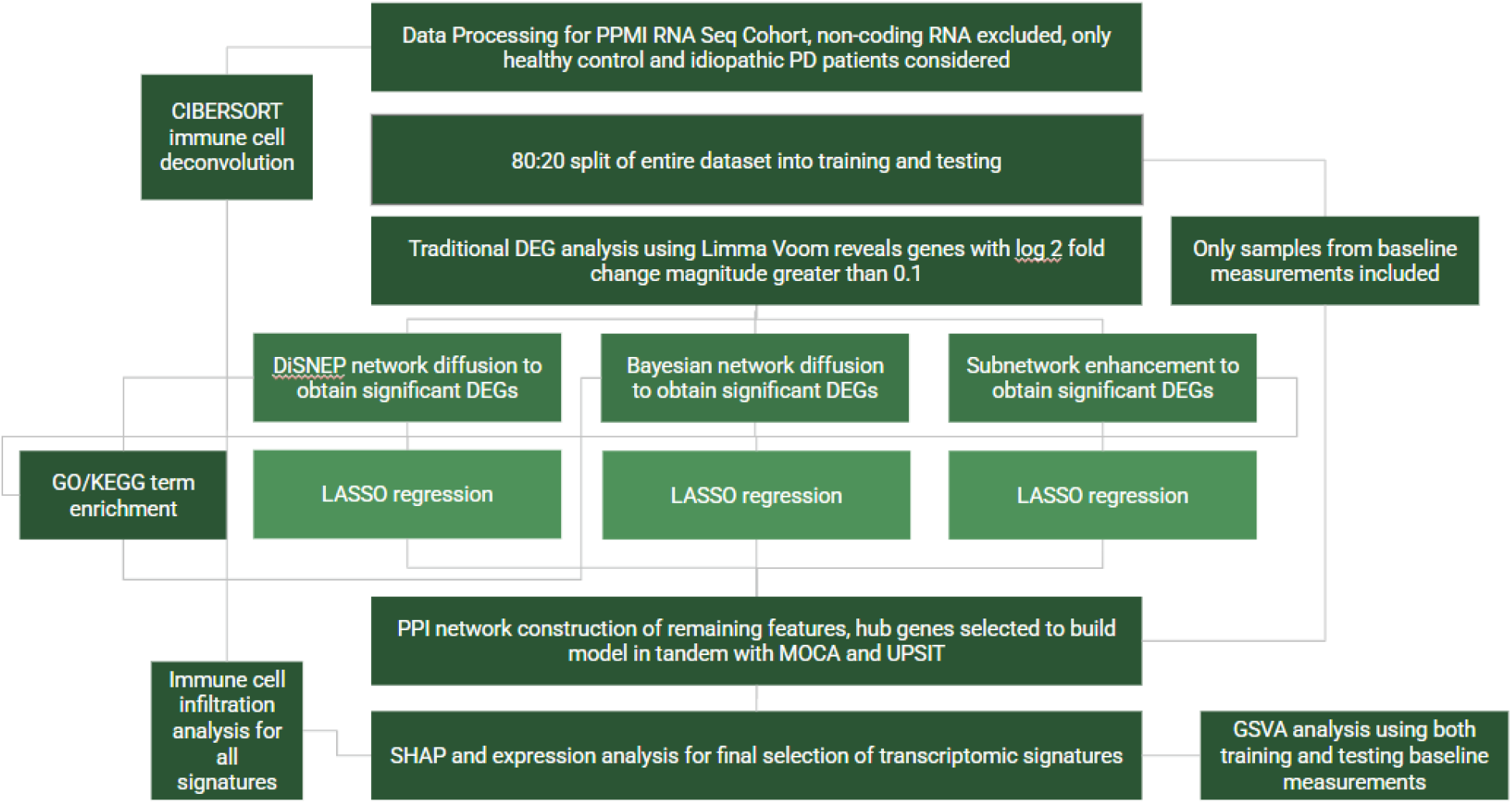
Flowchart of the statistical analysis strategy along with biological characterization.

## Results

### DiSNEP Network vs Bayesian Network Topology

Approximately 10% of all possible interactions between two genes were identified by the DiSNEP network as significant. However, only 0.03% of interactions between two genes were deemed existent by the Bayesian framework. To determine which network reflected the underlying biology more accurately, a scale free topology check was performed for both methods (Fig. 2A-B).

**Figure 2.**
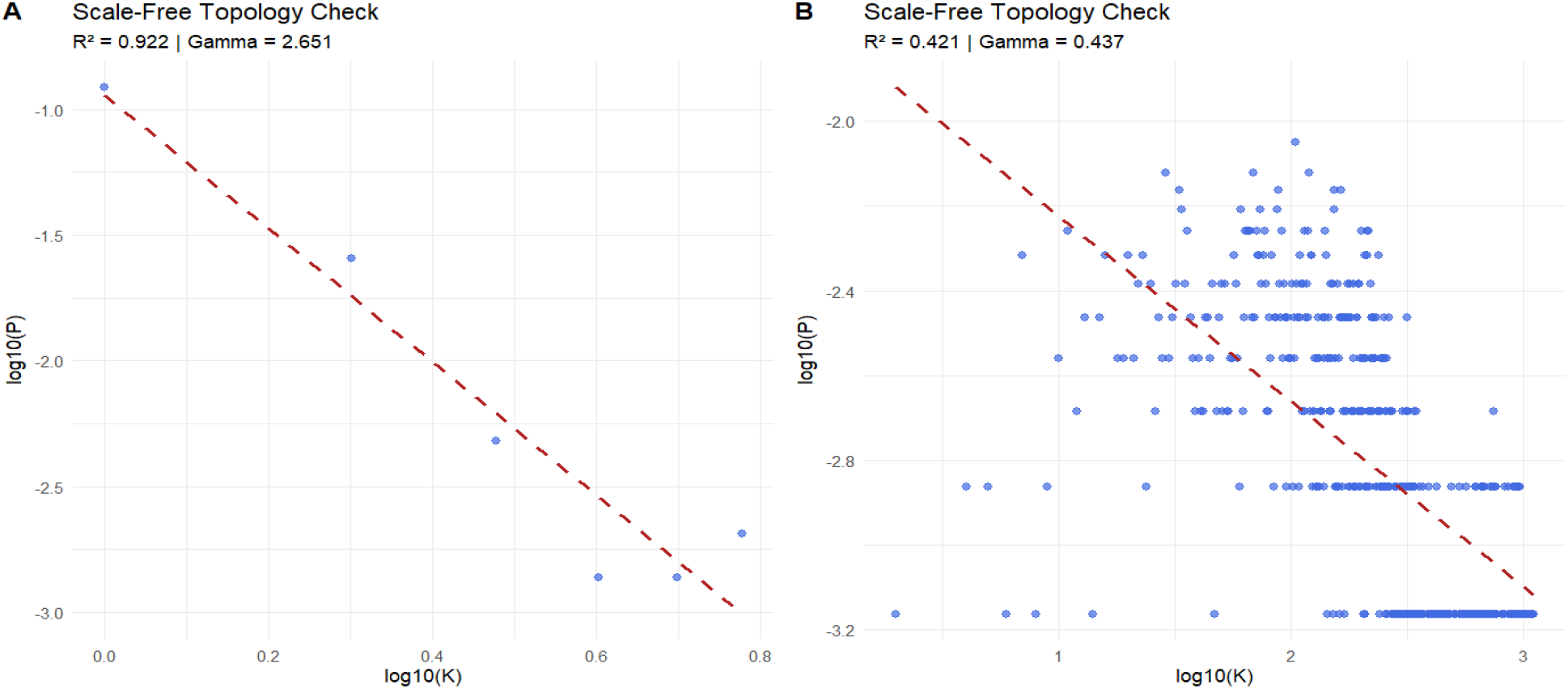
**(A)** Represents the topology check for the bayesian gene network. **(B)** Represents the topology check for the DiSNEP gene network.

From this analysis, not only does the Bayesian network have fewer connections, but also follows the power law, *log*(*P*) = *γ* * *log*(*k*), where k is the number of connections any gene has and P is the probability of genes with k number of connections. *γ* was fitted to be approximately 2.4. This means that the Bayesian network reflects the biological assumption that most connections between genes are governed by a few key hub genes. Meanwhile, the DiSNEP network breaks down in this assumption. The topology check also motivated the subnetwork analysis on the Bayesian network only instead of DiSNEP.

### GO/KEGG Enrichment Analysis and PPI Network

The GO analysis reveals significant enrichment in the biological processes module associated with cell chemotaxis, neutrophil mediated immunity, and peptide hormone processing (Fig. 3A). In terms of the cellular component module, these genes were enriched in tertiary granules and fincolin-1-rich granules. Both of these terms are associated with neutrophil activation, which is consistent with patterns in PD patients. The KEGG analysis revealed that the DEGs were enriched in Staphylococcus aureus infection [29], consistent with literature that validated the bacterium to produce amyloid phenol soluble modulins that accelerate protein aggregation, leading to neurodegeneration (Fig. 3B). Finally, after constructing a gene network from STRING, a total of 29 transcriptomic signatures were chosen. While the Bayesian-enhanced gene network was excellent for enhancing association signals, it was too sparse and conservative for selecting highly-connected genes. A total of 168 interactions were observed between 88 proteins with a p-value of 0.00014. Since the p-value of the network is less than 0.05, it can be concluded that these genes form biologically-relevant connections rather than forming a network based on random chance.

**Figure 3.**
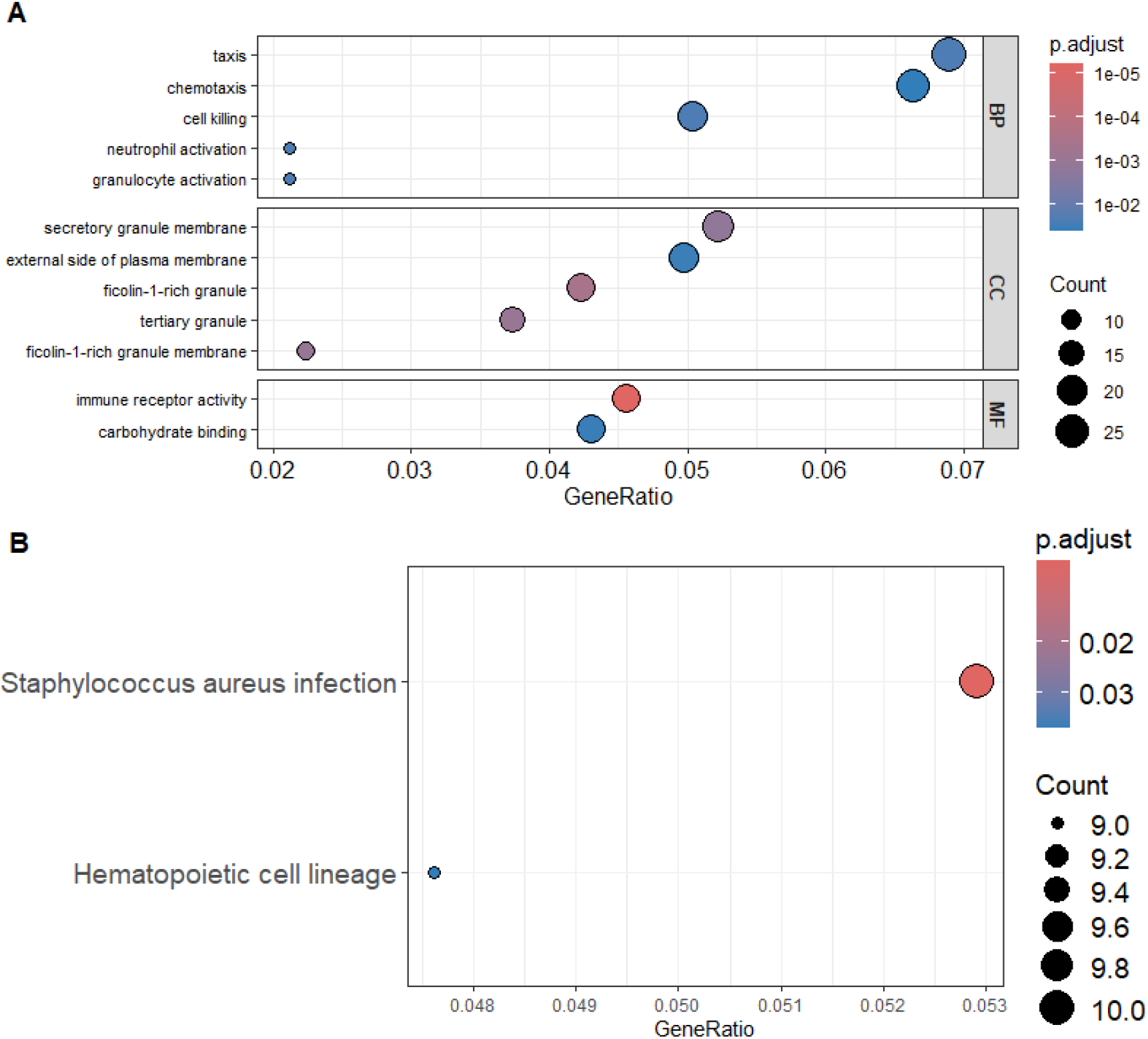
**(A)** GO terms where the significant network-recovered DEGs are significantly enriched in, sorted by biological processes (BP), cellular components (CC), and molecular function (MF). **(B)** KEGG terms that the network-recovered DEGs are significantly enriched in.

### Discrimination of iPD vs Healthy Control

Initially, the 29 hub genes–in tandem with UPSIT and MOCA–were able to achieve an AUC of 0.963 on the holdout test set with a random forest classifier. Using SHAP to narrow down candidates, the final 13 transcriptomic signatures that were chosen for optimization of classification were ALDH1A2, CA4, CD93, CCR9, ANXA3, IFI6, TREM1, IL1R2, CORIN, SIGLEC9, OCLN, PCSK6, and STK11. Combined with evaluations of UPSIT and MOCA scores, a new random forest classifier was trained and achieved a maximum test AUC of 0.972 (Fig. 4A). This exceeds the performance achieved from the panel with 29 signatures and also has the benefit of less genes to work with in a clinical application. Compared to the random forest model built only from UPSIT and MOCA scores, the hub genes provided a boost in AUC of approximately 0.054 (Fig. 4A). A Delong’s test was conducted to verify that the difference in performance of the two models is significant. Isolated, the 13 hub genes achieved a test AUC of 0.638 (Fig. 4A). When compared to the performance of other isolated signatures on the PPMI cohort, these genes still outperforms nearly every contemporary panel and matches the best performing panel without overfitting [10] [14]. At the point of highest overall accuracy, this test achieves a high specificity of 97% while achieving a sensitivity of 85.5% (Fig. 4B).

**Figure 4.**
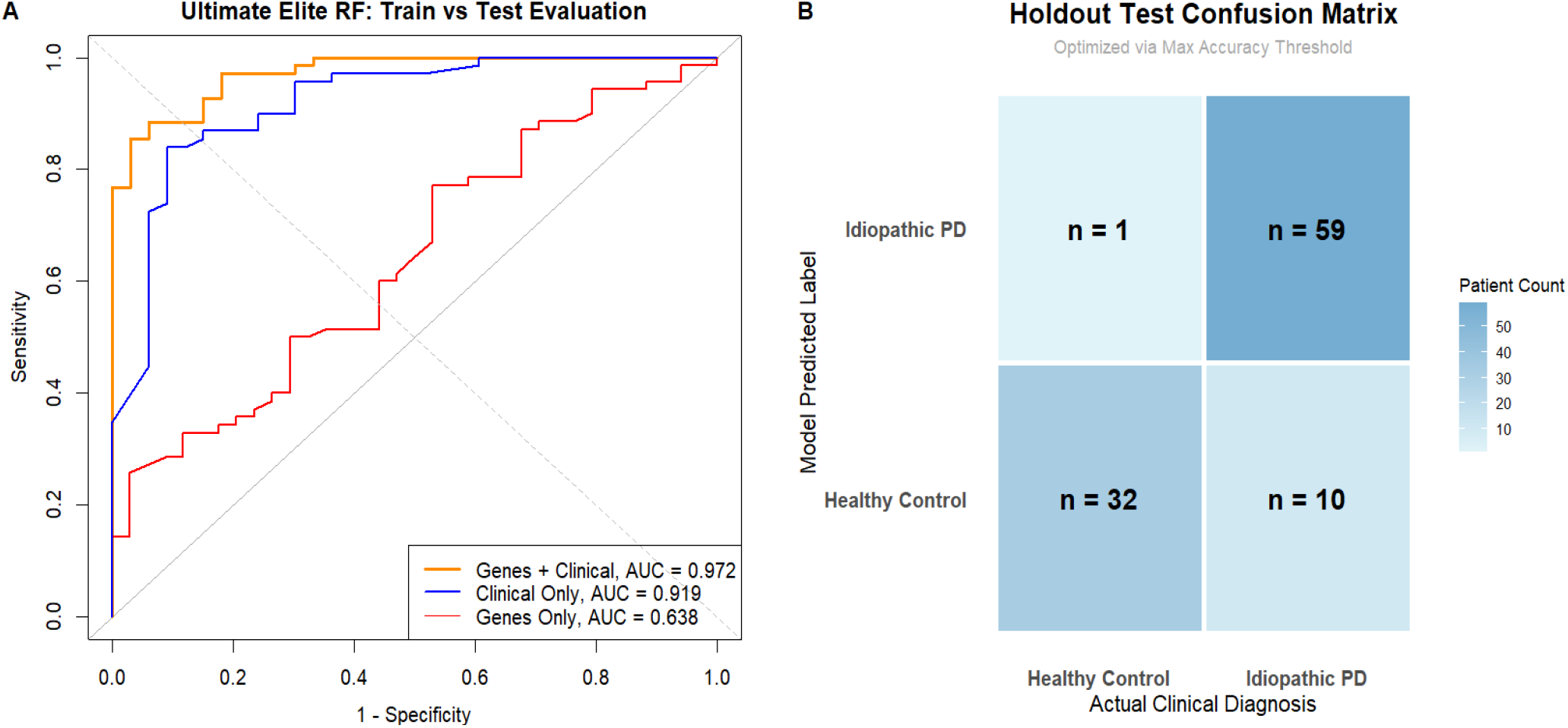
**(A)** Comparison of random forest model performance trained on transcriptomic signatures only vs clinical features only vs signatures + clinical features. **(B)** Confusion matrix that displays the sensitivity and analysis of the best performing model at the highest accuracy.

### Gene Expression in PD Pathways

By performing GSVA pathway enrichment analysis, many activated pathways were uncovered between high expression and low expression of each gene. Specifically, the upregulation of PCSK6, ALDH1A2, CA4, TREM1, IL1R2, CD93, and ANXA3 is positively associated with a coordinated shift in leukocyte transendothelial migration, chemokine signaling, and negative associated with immunodeficiency (Fig. 5). IFI6, STK11, TREM1 and ANXA3 are also of note due to their upregulation being positively associated with the PD pathway in the substantia nigra itself. Meanwhile, the upregulation of CCR9 is negatively associated with nearly all immune-response related terms while being positively associated with primary immunodeficiency. However, this is perfectly consistent with the expression patterns observed. The same trend is observed for OCLN, although its upregulation is positively associated with immunodeficiency, which contradicts the expected behavior. Lastly, SIGLEC9 displayed the strongest association with pathways involving leukocyte transendothelial migration, chemokine signaling cascade, toll-like receptor signaling and adipocyte signaling pathways (Fig. 5). Overall, the GSVA analysis yields great insight into the biological processes reflected by the 13 transcriptomic signatures and how they contribute to enhanced diagnosis.

**Figure 5.**
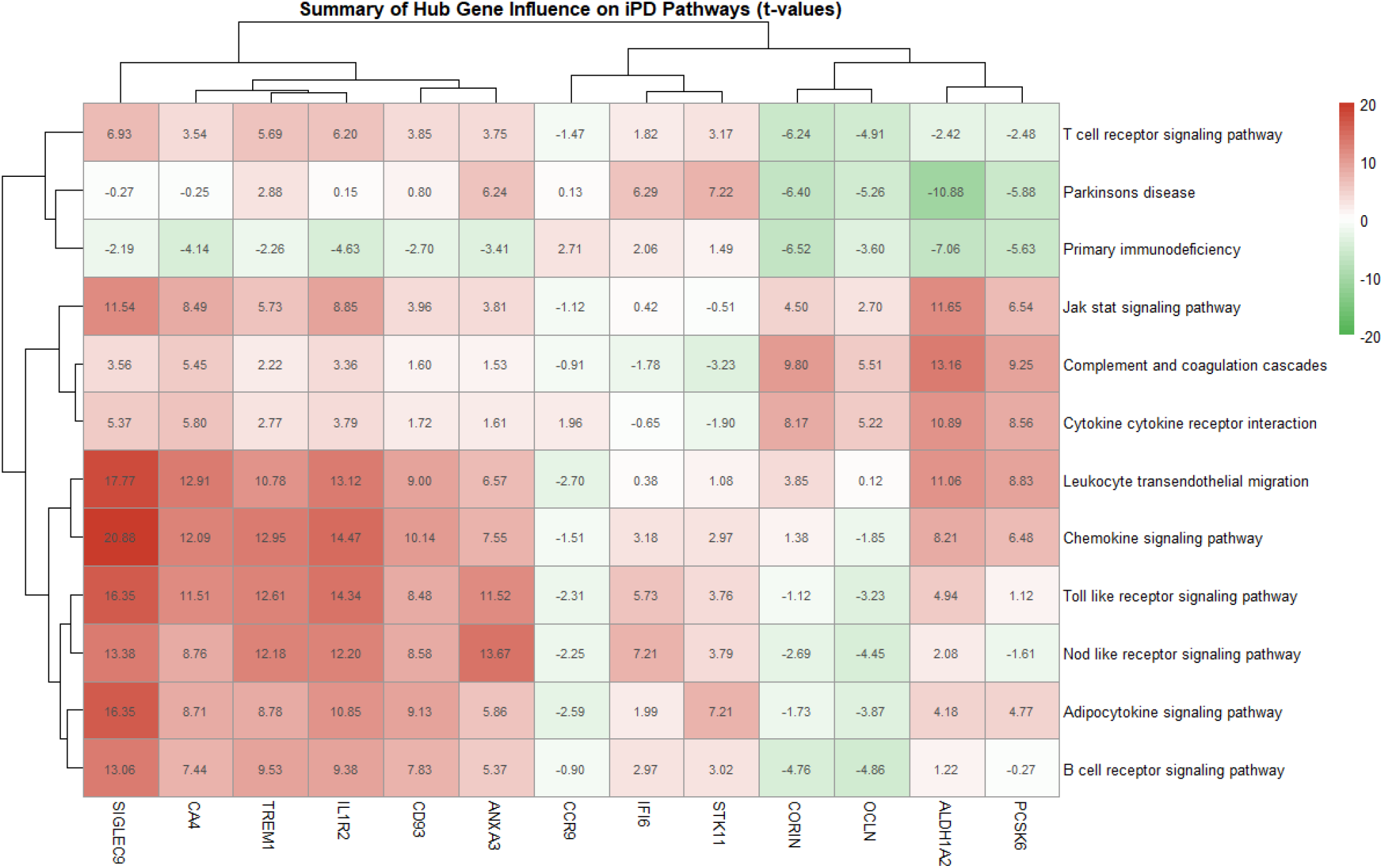
Correlation heatmap that demonstrates associations between pathways and all 13 signatures. Red signifies pathway is enriched with high expression of gene, while green signifies pathway is enriched with low expression of gene. t > 1.0 is a significant association.

### Immune Cell Infiltration

From the deconvolution, neutrophils and plasma cells were found to be significantly upregulated in iPD patients compared to HC patients, while naive CD4+ T cells were found to be significantly downregulated (Fig. 6A). Correlation analysis also illustrated that while the IFI6, OCLN, and CCR9 genes did not have any significant correlation with any significant cell types, the other 10 hub genes were all significantly associated with immune cell proportion changes (Fig. 6B). 6 of these genes were found to be positively correlated with neutrophils and plasma cells while being negatively correlated with naive CD4+ T cells. These patterns are consistent with the expression patterns of the hub genes. Specifically, ANXA3 and CA4 were found to have not only the most significant association with neutrophil proportions, but also the strongest correlations with neutrophils (Fig. 6B). It is important to note that STK11 has a negative yet significant correlation with neutrophil proportions, which is consistent with its observed downregulation. These results support that the underlying pathology captured by these genes focus on neuroinflammation.

**Figure 6.**
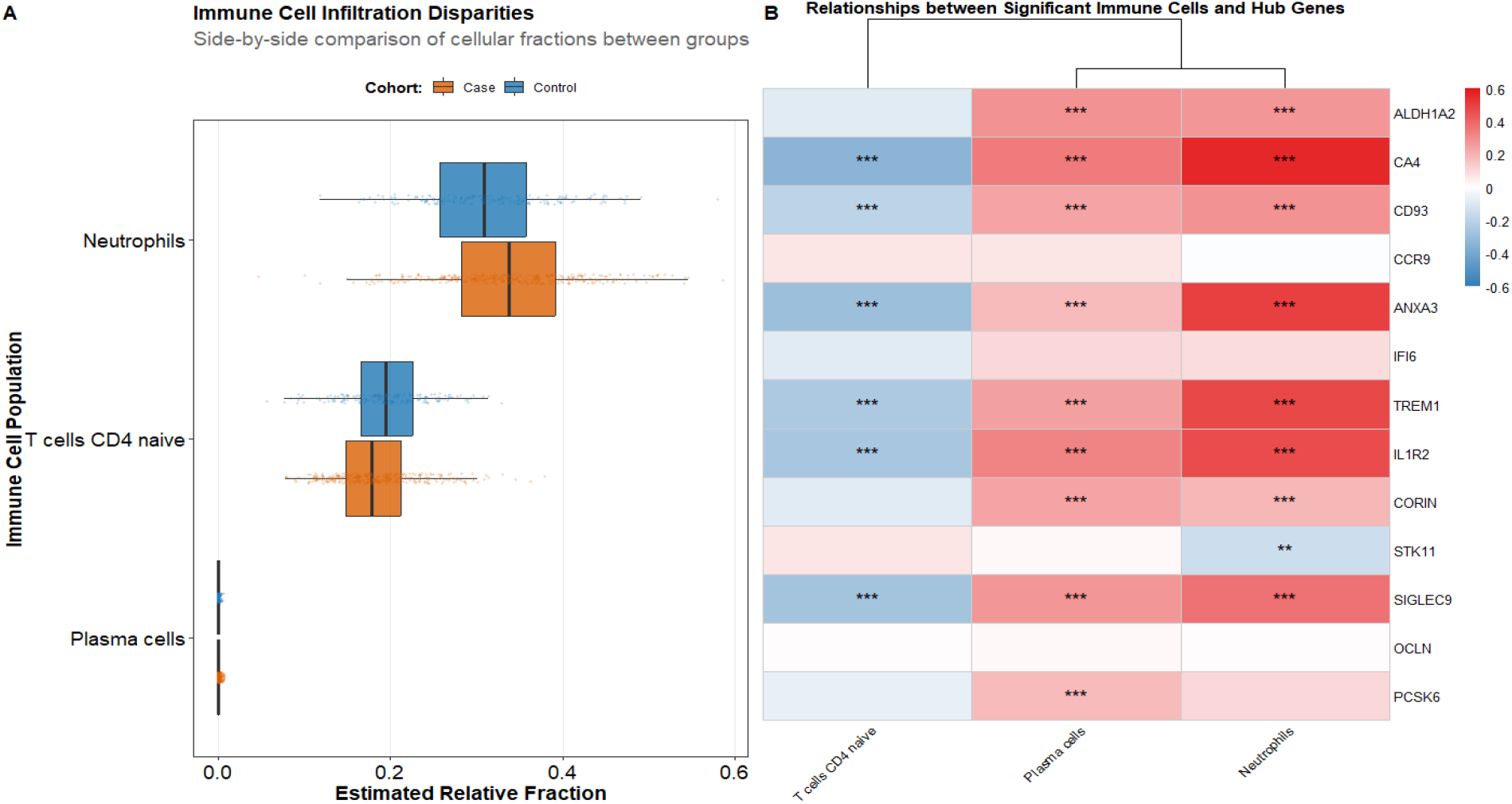
**(A)** Box plot of all immune cell types that are significantly different in proportions between healthy control and idiopathic PD patients. **(B)** Correlation analysis between the expression of signatures and the changes in cell-type proportions, */**/*** implies significance.

## Discussion

In this study, a novel panel of 13 transcriptomic signatures was developed that not only outperform most other panels, but display excellent discrimination when used alongside clinical evaluation. While almost none of these genes are novel discoveries, it is encouraging that all genes have been identified to be related to the progression of PD in one way or another.

Particularly, the genes selected all seem to be related to neuroinflammation and disruption of the blood brain barrier. The underlying biology contributes to the recovery of certain samples whose scores deviated from the average on MOCA and UPSIT tests. Specifically, several patients at baseline had MOCA scores that were comparable to the scores that healthy control patients at baseline had. Thus, a machine learning classifier trained only on those traits can easily miss these patients. Implementing a robust panel of transcriptomic signatures reveal signs of neuroinflammation that can relate to increased risk of PD progression. This can also be related to patients who were incorrectly identified as PD when they were in fact healthy. This patient had scores of MOCA and UPSIT that were comparable to idiopathic PD patients at baseline. It has been established that a low UPSIT score can translate to increased risk of PD progression.

However, it is also associated with other conditions such as COVID-19, other neurodegenerative disorders like Alzheimer’s Disease (AD), or just normal aging in general [30] [31]. The same principle can be applied to MOCA scores as well, only MOCA applies to an even wider range of conditions [32] [33] [34].

However, it is also important to note that not every patient with PD will exhibit excess inflammation, nor does inflammation indicate PD specifically rather than another neurodegenerative disease. This is shown by the signatures isolated only achieving limited performance in classification. Looking at the samples that were recovered by the UPSIT + MOCA model compared to the 13 signature model, the patients at baseline generally showed signs of neuroinflammation from their gene expression. Yet, this is not unique to PD progression, as neuroinflammation is a common marker of AD, multiple sclerosis (MS) or common infections that require a general immune response in the brain [35] [36]. This all demonstrates that neither the selected 13 genes nor the UPSIT/MOCA scores can be used as standalone tests in clinical practice. Combined together, however, each method corrects for the blindspots of the other and enhances general diagnostic accuracy. Furthermore, this approach can be applied to other complex diseases to boost classification performance. Complex diseases such as AD can also greatly benefit from early identification before emergence of symptoms. This method is not limited only to clinical traits, as proteomics and metabolics are other modes that have been demonstrated to work in tandem with blood transcriptomics.A recent example was the enhanced diagnosis of AD when using established plasma proteins complementary to gene expression [37]. Thus, more can be done to evaluate any additional data that can boost the current model’s performance.

Examining the genes individually, TREM 1 is already a well validated emerging biomarker in PD [38][39]. It acts as an amplifier of neuroinflammation in patients and promotes monocyte infiltration into the substantia nigra. The IFI6 gene has recently been identified to have an association with Parkinson’s disease. It functions primarily as an inflammation regulator. The upregulation of IFI6 is typically found in PD brain tissue from microglial activation [40]. It has also been identified to act locally in the mitochondria to control cell apoptosis, again suggesting a link with PD [41]. IL1R2 has also been identified to be a gene associated with PD, particularly in Southeast Asian populations [42][43]. While it functions to dampen the inflammatory process through competitive binding with Interleukin-1-β, this also acts as a signal that the body is trying to fight neuroinflammation [44]. The CORIN gene primarily functions to activate atrial natriuretic peptide (ANP) to control blood pressure [45]. It is also a surface marker to differentiate healthy dopamine-dopamine producing neuronal progenitors from induced pluripotent stem (iPS) cells in literature [46][47]. However, CORIN’s function as a gene in PD progression is not well researched yet. ALDH1A2 belongs to the ALDH1 family that is primarily localized in the cytoplasm of hepatocytes. While mainly responsible for acetaldehyde detoxification, studies have demonstrated that a dysregulation of the ALDH1A1 gene can lead to neuronal degradation due to build up of toxic metabolites [48][49]. However, it must be examined the mechanisms behind the upregulation of ALDH1A2 as identified in DEG analysis. CCR9, OCLN, CD93, and ANXA3 also have roles in the inflammatory process. Specifically, CCR9 is a gene heavily involved in T-cell differentiation [50]. It also acts as a cytokine receptor to guide effector T cells to the gut [51] [52]. Thus, its downregulation in PD patients is of interest. Meanwhile, OCLN functions to form tight junctions between endothelial and epithelial cells. Its downregulation has been associated with disruption of the blood brain barrier [53][54]. ANXA3 has been implicated in the neutrophil degranulation pathway. Such exacerbation can lead to tissue damage and neuronal damage in the context of PD [55]. This is consistent with the upregulation of the gene. CD93 promotes endothelial migration and also stimulates monocytes and macrophages [56] [57]. This is supported by the results from GSVA. The gene has also been validated as an emerging biomarker in recent PD research.

While CA4 is not as heavily involved in the immune response directly, literature has frequently shown that CA4 was highly expressed in PD patients and is a promising direction in novel targets for treatment [58]. Meanwhile, PCSK6 is a gene that functions as a key protease.

Specifically, inactive precursor proteins are cut by PCSK6 for the regulation of smooth muscle cells [59]. This is mainly to regulate the structure of cartilage and blood vessel walls. There is also literature that identifies PCSK6 as a potential risk gene in PD due to its duplication in siblings with PD compared to their healthy siblings [60]. In mice, it was also discovered that PCSK6–when overexpressed–can cause dopaminergic neurons to be vulnerable to degeneration [61]. SIGLEC9 belongs to the SIGLEC family. While the gene is often responsible as an inhibitor to inflammatory processes, the family is validated extensively in literature for its role in neurodegenerative diseases [62][63]. Thus, SIGLEC9 could be another brake gene that attempts to dampen the immune response. Finally, the STK11 gene encodes the LKB1 enzyme. The deletion of the LKB1 gene contributes to axon degeneration and motor dysfunction [64], which would be consistent with a downregulation of STK11 in PD patients. Overall, the genes in the panel capture a consistent narrative that enhances the diagnosis of PD without the observance of motor symptoms.

Almost all the genes in the 13 signatures show strong potential as targets for PD treatment. Furthermore, aside from CORIN, TREM1, and CCR9, almost all the other signatures were obtained from the LASSO regression using significant DEGs from Bayesian network analysis. Among those genes, PCSK6 was a gene that was not recovered by either traditional DEG analysis or DiSNEP gene network enhancement. While none of the genes identified from the Bayesian subnetwork analysis was included in the final panel, the module framework was able to identify ANPEP as a notable feature gene. Recent literature has identified ANPEP as being upregulated and exacerbating neuroinflammation in patients with PD [65] [66]. Thus, these results indicate that a Bayesian framework to gene interaction network and subnetwork analysis is a promising direction for future biomarker discovery.

Reverse-transcriptase qPCR (RT-qPCR) is a promising direction in adapting transcriptomic discovery into applicable clinical utility. With that in mind, it is important to know that certain genes that were identified to be DEGs may not be significant or even have different directions of regulation when applied to an external qPCR cohort. Furthermore, due to the heterogeneity of both patients and reaction conditions, it would be advantageous to select the valid genes and obtain the relative expression of said signatures against a set of housekeeping genes. From literature, genes such as GAPDH or PGK1 have been selected to have stable expression, with PGK1 often being used as a reference gene in liquid biopsies [67] [68]. However, it must be taken into account that stable gene expression often varies across varying conditions. Thus, the best procedure is to conduct RT-qPCR on candidate housekeeping genes. Then, using algorithms like geNorm or NormFinder, researchers can determine the most stable set as reference. Unlike the PPMI cohort dataset, the 13 transcriptomic signatures–assuming that all 13 are valid in an external RT-qPCR cohort–will be measured in ΔCt, representing the difference in the PCR cycle numbers between the genes of interest and reference genes. Finally, a new classification model would be built on the RT-qPCR data–in tandem with UPSIT and MOCA scores–followed by independent validation.

In regard to target specificity and sensitivity of a clinical test, it is best to use a liquid biopsy test as a screening for further confirmatory testing like DaTscan rather than a standalone tool. This way, the cost of a false positive can be around 2,500-5,000 USD for follow-up [69]. Furthermore, the economic burden on a patient for a RT-qPCR can reach as little as around 150 USD as a median cost [70]. For reference, a study demonstrated that clinical evaluation of patients with non-motor symptoms can take 3 years to receive a formal diagnosis on average [71]. Another study showed that the relative economic burden of treatment and therapy for a patient diagnosed with PD accumulates to around 28,000 USD to 30,000 USD in the first 3 years [72]. In this context, the cost of a false negative is relatively high compared to the cost of a false positive, encouraging high sensitivity alongside high specificity.

Although the current classification model would not be appropriate for clinical use, derivation of the cost approach for ROC curves was used to identify the optimal cutoff threshold as the point where the slope of the curve was approximately 0.99 [73]. The formula for the cost rule is shown below, where CFN is the cost of a false negative and CFP is the cost of a false positive and Pr is the prevalence of PD.

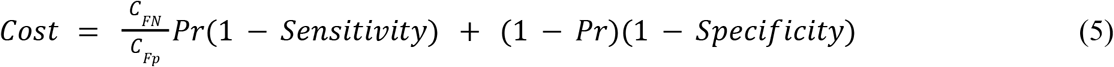

The prevalence of PD was determined to be approximately 1% for adults aged 60-65 [74]. The CFN and CFP were already determined to be 35,000 USD and 2,500 USD, respectively. Finally, the closest point on the ROC curve that achieved a slope of 0.99 achieved a sensitivity of 86.9% at a specificity of 93.9%, with an overall accuracy of 89.2%. Thus, this further demonstrates the potential of these 13 genes to be adopted in a clinical setting for effective early identification of idiopathic PD before emergence of major motor symptoms.

The key limitation of this study is that while the model achieves excellent discrimination in the holdout test set, it has not yet been validated in an external cohort. Thus, further analysis must be performed to examine how well these 13 signatures generalize in a different population like the Parkinson’s Disease Biomarkers Program (PDBP) cohort. Another limitation from this study is that the potential use of noncoding RNA (ncRNAs) signatures was not utilized or explored.

Specifically, many studies have focused on the potential of micro-RNAs (miRNAs) to enhance multi-omics classification of PD [75] [76] [77]. However, not only are these molecules low in abundance compared to long-coding RNAs, but miRNAs in particular have demonstrated low reproducibility when applied across multiple cohorts [78]. These factors limit the use of ncRNAs in clinical diagnosis. Nonetheless, applying network diffusion methods to identify ncRNAs of interest is a promising direction in the future. Finally, while the selected signatures are reflective of biological variation between idiopathic PD and healthy patients, their generalizability to PD patients with some form of genetic mutation remain yet to be seen. This study focuses solely on idiopathic PD patients due to difficulties in biomarker discovery with the lack of a single genetic cause.

## Conclusion

In conclusion, 13 transcriptomic signatures were observed to achieve excellent discrimination alongside clinical evaluation on the PPMI cohort. These 13 genes are implicated in immune-cell infiltration and neuroinflammation. Many of these genes were also recovered by diffusion through a Bayesian-constructed gene-interaction network. Finally, it is demonstrated that the combination of gene expression and clinical evaluation can boost predictive performance of PD diagnosis before emergence of motor symptoms. These findings provide insights into future examination of a clinically applicable test for the diagnosis of PD for effective intervention.

## Notes

https://fairsharing.org/

### Competing Interest Statement

The authors have declared no competing interest.

